# A Multi-Scale Computational Model of Excitotoxic Loss of Dopaminergic Cells in Parkinson’s Disease

**DOI:** 10.1101/2020.02.20.957704

**Authors:** Vignayanandam R. Muddapu, V. Srinivasa Chakravarthy

## Abstract

Parkinson’s disease (PD) is a neurodegenerative disorder caused by loss of dopaminergic neurons in Substantia Nigra pars compacta (SNc). Although the exact cause of the cell death is not clear, the hypothesis that metabolic deficiency is a key facor has been gaining attention in the recent years. In the present study, we investigate this hypothesis using a multi-scale computational model of the subsystem of the basal ganglia comprising Subthalamic Nucleus (STN), Globus Pallidus externa (GPe) and SNc. The model is a multiscale model in that interactions among the three nuclei are simulated using more abstract Izhikevich neuron models, while the molecular pathways involved in cell death of SNc neurons are simulated in terms of detailed chemical kinetics. Simulation results obtained from the proposed model showed that energy deficiencies occurring at cellular and network levels could precipitate the excitotoxic loss of SNc neurons in PD. At the subcellular level, the models show how calcium elevation leads to apoptosis of SNc neurons. The therapeutic effects of several neuroprotective interventions are also simulated in the model. From neuroprotective studies, it was clear that glutamate inhibition and apoptotic signal blocker therapies were able to halt the progression of SNc cell loss when compared to other therapeutic interventions, which only slows down the progression of SNc cell loss.

## INTRODUCTION

Parkinson’s disease (PD) is predominantly considered as motor disorder, which affects more than 6 million people around the world (Chaudhuri et al., 2006). It is caused by the loss of dopaminergic neurons in substantia nigra pars compacta (SNc) situated in the midbrain region (Fu et al., 2018). The cardinal symptoms of PD, such as tremor, rigidity, bradykinesia, and postural instability (Goldman and Postuma, 2014) are thought to be considered as the first sign of PD pathogenesis. However, other symptoms such as anosmia (loss of smell) (Omori and Okutani, 2019), constipation (Lubomski et al., 2019), sleep disorders (specifically rapid eye movement behavior sleep disorder) (Postuma et al., 2019), and depression (Bayram et al., 2019) also emerge well before motor impairments. Recently, several studies pointed to the fact that even before emergence of motor and non-motor symptoms, pathogenesis begins with metabolic abnormalities occurring at different levels of neural hierarchy: subcellular, cellular and network levels (Bolam and Pissadaki, 2012; Limphaibool et al., 2018; Muddapu et al., 2019; Nam et al., 2018; Pacelli et al., 2015; Pissadaki and Bolam, 2013). With the help of a computational model, Muddapu et al (Muddapu et al., 2019) have recently suggested that the excitotoxic loss of SNc cells might be due to energy deficiencies occurring at different levels in the neural hierarchy.

If metabolic factors are indeed the deep underlying reasons behind PD pathogenesis, it is a hypothesis that deserves closer attention because any positive proof regarding the role of metabolic factors puts an entirely new spin on PD research. Unlike current therapeutic approaches that manage the symptoms rather than provide a cure, the new approach can, in principle, point to a more lasting solution. If inefficient energy delivery or energy transformation mechanisms are the reason behind degenerative cell death in PD, relieving the metabolic load on the vulnerable neuronal clusters, by intervening through surgical and/or pharmacological approaches could prove to be a decisive treatment for PD, a disease that previously proved itself to be intractable to current therapeutic approaches.

In this work, using computational models, we investigate the hypothesis that the major cause of SNc cell loss in PD might be due to energy deficiency occurring in SNc neurons. In the proposed modeling study, we focus on excitotoxicity in SNc caused by the subthalamic nucleus (STN), which is precipitated by energy deficiency and understand the mechanism behind neurodegeneration during excitotoxicity. Moreover, it aims to suggest therapeutic interventions that can reduce the metabolic burden on the SNc neurons, which in turn delay the progression of SNc cell loss in PD. The previous model (Muddapu et al., 2019; Muddapu and Chakravarthy, 2017) uses a slightly abstract single neuron model of SNc with simplified representation of programmed cell death and explores the network level interactions that possibly lead to cell death. In the present paper we use a biophysically elaborate model of SNc neurons and study the link between energy deficiency and molecular level changes that occur inside SNc neurons under PD conditions. Specifically, we consider intracellular molecular mechanisms such as energy metabolism, dopamine turnover processes, calcium buffering mechanisms, and apoptotic signal pathways. Using such a hybrid model, we were able to show excitotoxic loss of SNc neurons, which was precipitated by energy deficiency and postulated possible mechanism behind excitotoxic neurodegeneration. Moreover, we also explore various therapeutic interventions to halt or slow down the progression of SNc cell loss.

## METHODS

The proposed hybrid excitotoxicity model (HEM) in PD consists of three nuclei from basal ganglia, namely STN, SNc, and globus pallidus externa (GPe). SNc neuron was modeled as a biophysical neuronal model, and STN and GPe neurons (Muddapu et al., 2019) are modeled using the Izhikevich neuron model (Izhikevich, 2007). Neurons in each nucleus are arranged as a two-dimensional lattice (Figure 1). All the simulations were carried on MATLAB (RRID: SCR_001622) platform where all models numerically integrated with a time step of 0.1 *ms*.

**Figure 1:**
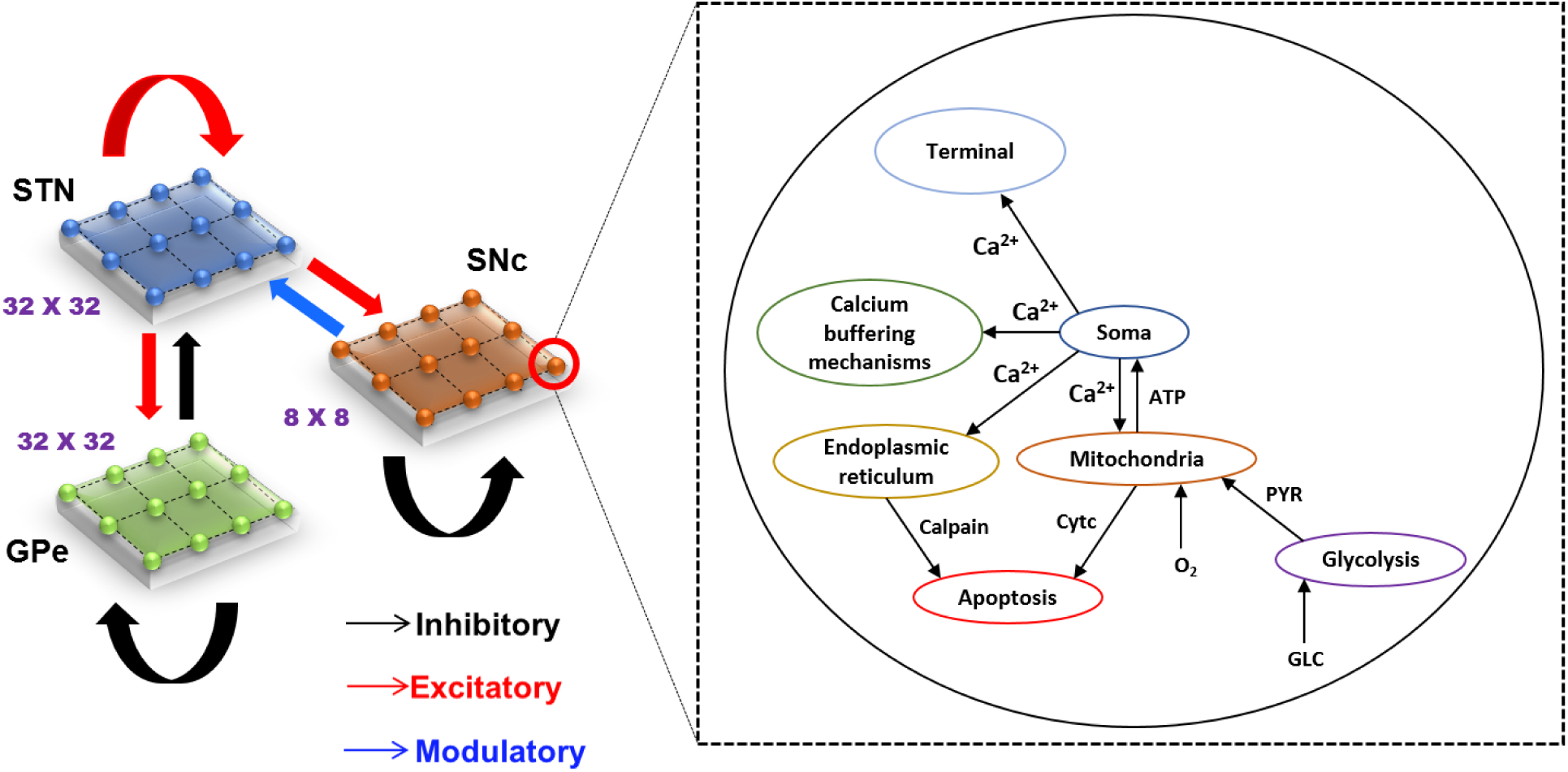
Model architecture of excitotoxicity In SNc. SNc, substantia nigra pars compacta; STN, subthalamic nucleus; GPe, globus pallidus externa; Ca^2+^, calcium; ATP, adenosine triphosphate; Cytc, cytochrome c; PYR, pyruvate; O_2_, oxygen; GLC, glucose.

### Izhikevich (Spiking) Neuronal Model (STN, GPe)

The Izhikevich neuronal models are capable of exhibiting biologically realistic firing patterns, at a relatively low computational expense (Izhikevich, 2003). The proposed model of HEM consists of GPe and STN neurons are modeled as Izhikevich spiking neuron model, where the Izhikevich parameters are adopted from the literature (Mandali et al., 2015; Michmizos and Nikita, 2011). Based on the anatomical data of rat basal ganglia, the neuronal population sizes in the model are selected (Arbuthnott and Wickens, 2007; Oorschot, 1996). The external bias current (*I*^*x*^) was adjusted to match the firing rate of nuclei with published data (Tripathy et al., 2015).

The Izhikevich neuron model of GPe and STN consists of two variables namely membrane potential (*v*^*x*^), and membrane recovery variable (*u*^*x*^):

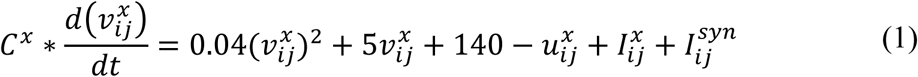

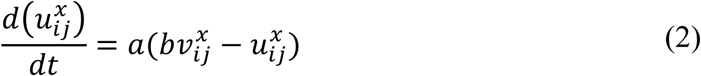

Resetting:

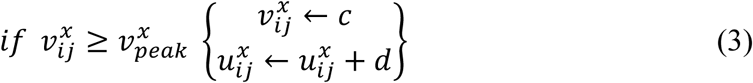

where, 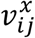, and 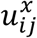 are the membrane potential, and the membrane recovery variables of a neuronal type *x* at the location (*i, j*) respectively, 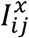, and 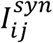 are the external bias, and total synaptic currents received to a neuronal type *x* at the location (*i, j*) respectively, *C*^*x*^ is the membrane capacitance of a neuronal type *x*, {*a, b, c, d*} are Izhikevich parameters, 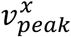 is the maximum (peak) membrane voltage set to a neuronal type *x* (where *x* = *STN or GPe*).

### Biophysical (Conductance-Based) Neuron Model (SNc)

The biophysical neuron model of SNc in the proposed HEM was adapted from (Muddapu and Chakravarthy, 2020). The detailed biophysical model of SNc neuron consists of cellular and molecular processes such as ion channels (active ion pumps, ion exchangers), calcium buffering mechanisms (calcium-binding proteins, organelles sequestration of calcium), energy metabolism pathways (glycolysis and oxidative phosphorylation), dopamine turnover processes (synthesis, storage, release, reuptake and metabolism), apoptosis (endoplasmic reticulum-stress and mitochondrial-induced apoptosis). The dynamics of SNc membrane potential (*v*^*SNc*^) is given as,

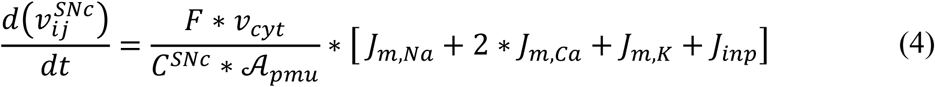

where *F* is the Faraday’s constant, *C*^*SNc*^ is the SNc membrane capacitance, *v*_*cyt*_ is the cytosolic volume, 𝒜_*pmu*_ is the cytosolic area, *J*_*m,Na*_ is the sodium membrane ion flux, *J*_*m,Ca*_ is the calcium membrane ion flux, *J*_*m,K*_ is the potassium membrane ion flux, *J*_*inp*_ is the overall input current flux. A more detailed description of the SNc neuron model was provided in (Muddapu and Chakravarthy, 2020).

In the proposed model, intracellular calcium concentration in the SNc neuron is dependent on calcium-binding proteins, mitochondria (MT) and endoplasmic reticulum (ER) (Zaichick et al., 2017; Zündorf and Reiser, 2011). The intracellular calcium concentration dynamics ([*Ca*_*i*_]) of the SNc neuron (Muddapu and Chakravarthy, 2020) is given by,

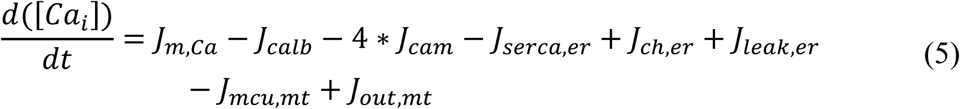

where, *J*_*m,Ca*_ is the flux of calcium ion channels, *J*_*calb*_ is the calcium buffering flux by calbindin, *J*_*cam*_ is the calcium buffering flux by calmodulin, *J*_*serca,er*_ is the calcium buffering flux by ER uptake of calcium through sarco/endoplasmic reticulum calcium-ATPase, *J*_*ch,er*_ is the calcium efflux from ER by calcium-induced calcium release mechanism, *J*_*leak,er*_ is the calcium leak flux from ER, *J*_*mcu,mt*_ is the calcium buffering flux by MT uptake of calcium through mitochondrial calcium uniporters (MCUs), *J*_*out,mt*_ is the calcium efflux from MT through sodium-calcium exchangers, mitochondrial permeability transition pores (PTPs), and non-specific leak flux.

### Synaptic Connections

The synaptic connectivity among different neuronal populations was modeled as a standard single exponential model of postsynaptic currents (Humphries et al., 2009) as follows:

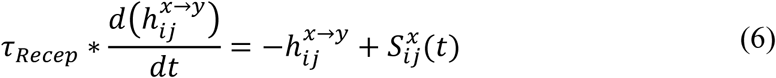

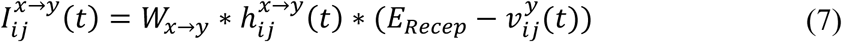

The N-Methyl-D-aspartic Acid (NMDA) current was regulated by voltage-dependent magnesium channels which were modeled as,

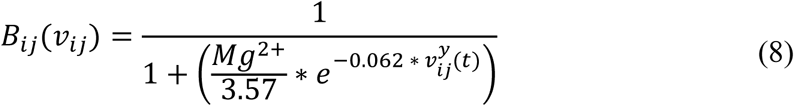

where, 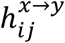 is the gating variable for the synaptic current from neuronal type *x* to neuronal type *y* (where *x* → *y* = {*STN* → *STN, STN* → *SNc, STN* → *GPe, GPe* → *STN, GPe* → *GPe*}), τ_*Recep*_ is the decay constant for the synaptic receptor, 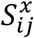 is the spiking activity of a neuronal type *x* at time *t, W*_*x*→*y*_ is the synaptic weight from neuron *x* to *y*, 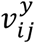 is the membrane potential of the neuronal type *y* at the location (*i, j*), *E*_*Recep*_ is the receptor-associated synaptic potential (*Recep* = GABA/AMPA/NMDA), and [*Mg*^2+^] is the magnesium ion concentration. The time constants of Gamma-Amino Butyric Acid (GABA), Alpha-amino-3-hydroxy-5-Methyl-4-isoxazole Propionic Acid (AMPA) and NMDA in STN and GPe were chosen from (Götz et al., 1997) are given in Table 1.

**Table 1:** Parameter values used in the proposed HEM.

| Parameter(s) | STN | GPe |
| --- | --- | --- |
| Izhikevich parameters |  |  |
| $a \text{ (ms}^{-1}\text{)},$ | $a = 0.005,$ | $a = 0.1,$ |
| $b \text{ (pA.mV}^{-1}\text{)},$ | $b = 0.265,$ | $b = 0.2,$ |
| $c \text{ (mV)},$ | $c = -65,$ | $c = -65,$ |

| $d$ (pA) | $d = 1.5$ | $d = 2$ |
| --- | --- | --- |
| External current ($I^x$) | 3 pA | 4.25 pA |
| Maximum peak of voltage ($v_{peak}^x$) | 30 mV | 30 mV |
| Membrane capacitance ($C^x$) | 1 $\mu F$ | 1 $\mu F$ |
| Number of laterals ($nlat^x$) | 11 | 15 |
| Radius of Gaussian laterals ($R^x$) | 1.4 | 1.6 |
| Synaptic strength within laterals ($A^x$) | 1.3 | 0.1 |
| Time decay constant for AMPA ($\tau_{AMPA}$) | 6 ms | 6 ms |
| Time decay constant for NMDA ($\tau_{NMDA}$) | 160 ms | 160 ms |
| Time decay constant for GABA ($\tau_{GABA}$) | 4 ms | 4 ms |
| Synaptic potential of AMPA receptor ($E_{AMPA}$) | 0 mV | 0 mV |
| Synaptic potential of NMDA receptor ($E_{NMDA}$) | 0 mV | 0 mV |
| Synaptic potential of GABA receptor ($E_{GABA}$) | -60 mV | -60 mV |
| Concentration of Magnesium ($Mg^{2+}$) | 1 mM | 1 mM |

### Lateral Connections

The lateral connections in SNc, STN and GPe, are modeled as Gaussian neighborhoods (Muddapu et al., 2019),

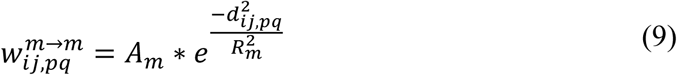

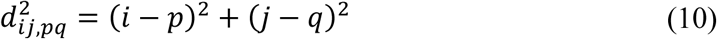

where, 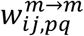 is the weight of lateral connection strength of a neuronal type *m* at the location (*i, j*), *d*_*ij,pq*_ is the distance of neuron at the location (*i, j*) from the center neuron (*p, q*), *R*_*m*_ is the standard deviation of Gaussian, *A*_*m*_ is the amplitude of lateral synaptic strength (where *m* = *GPe or STN or SNc*).

The lateral connections within SNc and GPe populations are considered as inhibitory and within STN as excitatory (Muddapu et al., 2019) (Figure 1). The lateral currents in the STN and GPe are modeled similar to equations (6-8) and in the case of SNc which was modeled as,

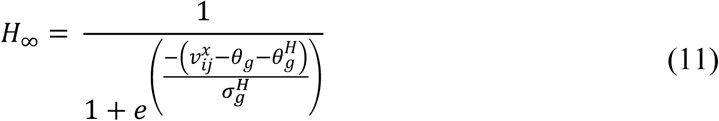

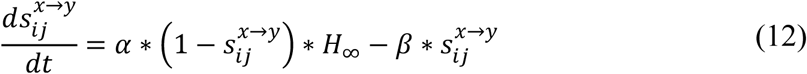

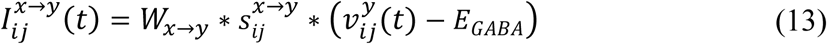

where 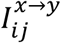 is the synaptic current from neuronal type *x* to neuronal type *y, W*_*x*→*y*_ is the weight of synaptic strength from neuronal type *x* to neuronal type *y*, 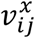 and 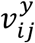 are the membrane potential of the neuronal type *x* and *y* respectively at the location (*i, j*), *E*_*GABA*_ is the GABAergic receptor reversal potential, 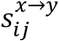 is the synaptic gating variable from neuronal type *x* and *y* at the location (*i, j*) (where *x* → *y* = {*SNc* → *SNc*} only). The parametric values of 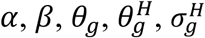 are adopted from (Rubin and Terman, 2004) and given in Table 2.

**Table 2:** Parameter values used for SNc lateral connections.

| Parameter | Value | Parameter | Value |
| --- | --- | --- | --- |
| Number of laterals ($nlat^x$) | 5 | $\theta_g$ | 20 mV |
| Radius of Gaussian laterals ($R^x$) | 1.6 | $\theta_g^H$ | -57 mV |
| Synaptic strength within laterals ($A^x$) | 0.1 | $\sigma_g^H$ | 2 mV |
| Synaptic conductance ($W_{x \rightarrow y}$) | 0.01 | $\alpha$ | 2 ms<sup>-1</sup> |
| Synaptic potential of GABA receptor ($E_{GABA}$) | 63.45 mV | $\beta$ | 0.08 ms<sup>-1</sup> |

### Effect of Dopamine on Synaptic Plasticity

Dopamine (DA) modulated lateral connection strength in STN, SNc, and GPe populations. As DA level increases, the lateral connection strength in GPe and SNc increases, whereas, in the case of STN, it decreases. DA-modulation of lateral connection strength is modeled as,

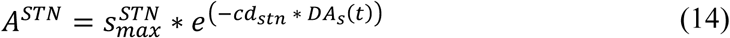

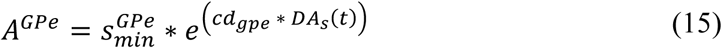

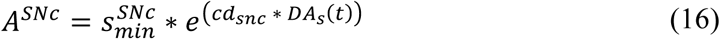

where, 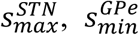, and 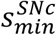 are lateral connection strengths of *STN, GPe*, and *SNc*, respectively where dopamine effect on the basal spontaneous activity of the neuronal population was minimal, *cd*_*stn*_, *cd*_*gpe*_, and *cd*_*snc*_ are the parameter values which modulate the influence of dopamine on the lateral connections in *STN, GPe*, and *SNc* populations respectively, *DA*_*s*_(*t*) is the instantaneous DA level. The instantaneous DA was estimated by the spatial average DA concentration of all the terminals at a given instant. All parameter values are given in Table 3.

**Table 3:** Parameter values of dopamine effect on target neuronal populations.

| Parameter | Value | Parameter | Value |
| --- | --- | --- | --- |
| $s_{max}^{STN}$ | 1.3 | $cd_{stn}$ | 4.87 |
| $s_{min}^{GPe}$ | 0.1 | $cd_{gpe}$ | 7 |
| $s_{min}^{SNc}$ | $1 \times 10^{-6}$ | $cd_{snc}$ | 4.6055 |
| $cd2$ | 0.1 | $w_{STN \rightarrow SNc}$ | 0.3 |
| $w_{GPe \rightarrow GPe}$ | 1 | $w_{SNc \rightarrow SNc}$ | 0.01 |
| $w_{STN \rightarrow GPe}$ | 1 | $w_{GPe \rightarrow STN}$ | 20 |
| $w_{STN \rightarrow STN}$ | 1 | $F_{STN \rightarrow SNc}$ | $1 \times 10^{-5}$ |

The post-synaptic effects of DA in SNc, STN and GPe are modeled similar to (Muddapu et al., 2019),

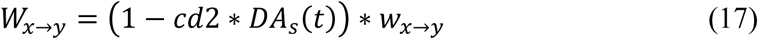

where, *w*_*x*→*y*_ is the synaptic weight (*STN* → *GPe, GPe* → *STN, STN* → *STN, GPe* → *GPe, STN* → *SNc, SNc* → *SNc*), *cd*2 is the parameter that affects the post-synaptic current, *DA*_*s*_(*t*) is the instantaneous dopamine level.

### Total Synaptic Current Received by Each Neuronal Type

#### SNc

The total synaptic current received by a *SNc* neuron at the lattice position (*i, j*) is the summation of the glutamatergic input from the *STN* neurons, considering both *NMDA* and *AMPA* receptor activation, and lateral GABAergic current from other *SNc* neurons.

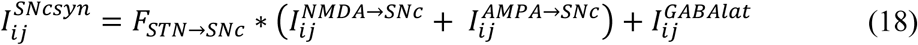

where, 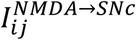 and 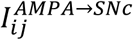 are the glutamatergic currents from the *STN* neurons corresponding to *NMDA* and *AMPA* receptors activation respectively; 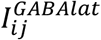 is the lateral GABAergic current from other *SNc* neurons; *F*_*STN*→*SNc*_ is the scaling factor for the glutamatergic current from *STN* neuron.

#### GPe

The total synaptic current received by a *GPe* neuron at the lattice position (*i, j*) is the summation of the glutamatergic input from the *STN* neurons considering both *NMDA* and *AMPA* receptors activation and the lateral GABAergic current from other *GPe* neurons.

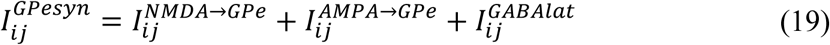

where, 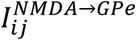 and 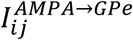 are the glutamatergic currents from *STN* neuron considering both *NMDA* and *AMPA* receptors activation respectively; 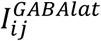 is the lateral GABAergic current from other *GPe* neurons.

#### STN

The total synaptic current received by a *STN* neuron at the lattice position (*i, j*) is the summation of the GABAergic input from the *GPe* neurons and the lateral glutamatergic input from other *STN* neurons considering both *NMDA* and *AMPA* receptors activation.

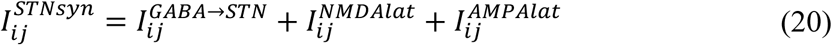

where, 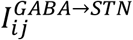 is the GABAergic current from *GPe* neuron; 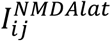 and 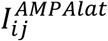 are the lateral glutamatergic currents from other *STN* neurons considering both *NMDA* and *AMPA* receptors activation respectively.

### Calcium-Induced Neurodegeneration in SNc

Calcium plays a dual role in living organisms – as a survival factor or a ruthless killer (Orrenius et al., 2003). For the survival of neurons, minimal (physiological) levels of glutamate stimulation are required. Under normal conditions, calcium concentration within a cell is tightly regulated by pumps, transporters, calcium-binding proteins, ER and MT (Surmeier et al., 2011; Wojda et al., 2008). Due to prolonged calcium influx driven by excitotoxicity, calcium homeostasis within the cell is disrupted, which results in cellular imbalance, leading to activation of apoptotic pathways (Bano and Ankarcrona, 2018). The SNc soma undergoes degeneration when there is an abnormal calcium build up inside the cell resulting in calcium loading inside ER and MT, which leads to ER stress-induced and MT-induced apoptosis respectively (Malhotra and Kaufman, 2011). The proposed model will also include the apoptotic processes inside SNc neuron, which get activated when calcium levels in the neuron cross a certain threshold as a result of overexcitation and/or metabolic deficiency. In the proposed HEM model, we incorporate a mechanism of programmed cell death, whereby an SNc neuron under high stress (high calcium levels) kills itself. The stress in a given SNc neuron is observed by monitoring the intracellular calcium concentrations in the cytoplasm, ER, and MT.

The SNc neuron undergoes ER-stress-induced apoptosis when calcium levels in ER cross a certain threshold (*ER*_*thres*_). Under such conditions, the particular SNc neuron gets eliminated as follows,

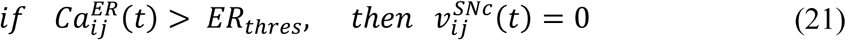

where, 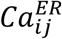 is the calcium concentration in the ER, *ER*_*thres*_ is the calcium concentration threshold after which ER-stress induced apoptosis gets initiated (*ER*_*thres*_ = 2.15 ×10^−3^ *mM*), 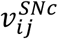 is the membrane voltage of neuron at the lattice position (*i, j*).

The SNc neuron undergoes mitochondria-induced apoptosis when calcium levels in mitochondria cross a certain threshold (*MT*_*thres*_). Then that particular SNc neuron will be eliminated as follows,

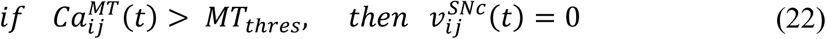

where, 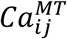 is the calcium concentration in mitochondria, *MT*_*thres*_ is the calcium concentration threshold after which mitochondria-induced apoptosis gets initiated (*MT*_*thres*_ = 0.0215 *mM*), 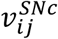 is the membrane voltage of neuron at the lattice position (*i, j*).

### Neuroprotective Strategies

Any therapeutic interventions which result in slow down of SNc cell loss can be considered as neuroprotective in nature. It can be achieved either by blockage of glutamatergic receptors on SNc or by attenuate the pathological oscillations in STN pharmacologically or surgically (Rodriguez et al., 1998). Just as in the previous model (Muddapu et al., 2019), glutamate inhibition, dopamine restoration, subthalamotomy, and deep brain stimulation therapies were implemented in the proposed HEM model. In addition to these therapeutics, since the present model captures several molecular mechanisms at subcellular level, we simulate therapeutic interventions at cellular level such as calcium channel blockers, enhancement of calcium-binding proteins (CBPs) expression, and apoptotic signal blockers.

#### Glutamate Inhibition Therapy

In glutamate inhibition therapy (such as MK-801, NBQX, LY-404187), the excitatory drive from STN neurons to SNc neurons is reduced by altering synaptic weight from STN to SNc (*W*_*STN*→*SNc*_) (Johnson et al., 2009; Zhang et al., 2019). In the proposed excitotoxicity model, the glutamate inhibition therapy was implemented by the following criterion,

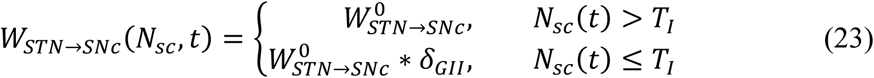

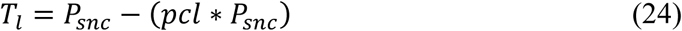

where, *W*_*STN*→*SNc*_(*N*_*sc*_, *t*) is the change in synaptic weight of STN to SNc connection based on the number of SNc cells surviving (*N*_*sc*_(*t*)) at a particular time (*t*), 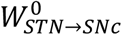 is the initial synaptic weight of STN to SNc connection, *N*_*sc*_(*t*) is the number of SNc neurons surviving at a particular time (*t*), *δ*_*GII*_ is the extent of glutamate inhibition, *pcl* is the percentage of SNc cell loss (25 %) at which intervention of a particular treatment was employed (*pcl* = 0.25), *P*_*snc*_ is the population size of SNc neurons, *T*_*l*_ represents the number of SNc cells surviving at which intervention of a particular treatment was employed. In the present study, the therapeutic intervention is given at 25% SNc cell loss.

#### Dopamine Restoration Therapy

In dopamine restoration therapy (such as levodopa, DA agonists), the dopamine drive to STN neurons is restored by increased by administration of external dopamine (*δ*_*DAA*_) (Vaarmann et al., 2013). In the proposed excitotoxicity model, the dopamine restoration therapy was implemented by the following criterion,

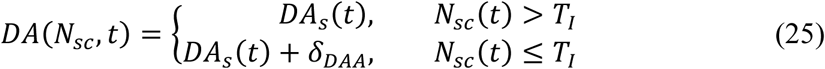

where, *DA*(*N*_*sc*_, *t*) is the change in dopamine level based on the number of SNc cells surviving at a particular time (*t*) (*N*_*sc*_(*t*)), *DA*_*s*_(*t*) is the instantaneous DA level, *N*_*sc*_(*t*) is the number of SNc neurons surviving at a particular time (*t*), *δ*_*DAA*_ is the extent of dopamine content restoration, *T*_*l*_ represents the number of SNc cells surviving at which intervention of a particular treatment was employed.

#### Subthalamotomy

In subthalamotomy therapy, the excitatory drive from STN neurons to SNc neurons is reduced by silencing or removing STN neurons (*δ*_*les*_) (Jourdain et al., 2014; Wallace et al., 2007). In the proposed excitotoxicity model, the subthalamotomy therapy was implemented by the following criterion,

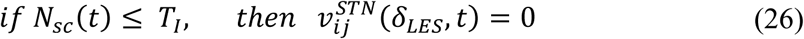

where, *δ*_*LES*_ is the percentage of STN lesioned which is in the following range: {5, 10, 20, 30, 40, 50, 60, 70, 80, 90, 100}, *N*_*sc*_(*t*) is the number of SNc neurons surviving at a particular time (*t*), *T*_*l*_ represents the number of SNc cells surviving at which intervention of a particular treatment was employed.

#### Deep Brain Stimulation (DBS) in STN

In our previous study (Muddapu et al., 2019), we proposed that the neuroprotective effect of DBS was due to increased axonal and synaptic failures in the stimulation site (Rosenbaum et al., 2014). Besides, we also suggested that biphasic current with four contact point (FCP) stimulation configuration showed a maximal neuroprotective effect. The DBS parameters such as amplitude (*A*_*DBS*_), frequency 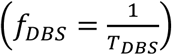, and pulse width (*δ*_*DBS*_) were adjusted by using clinical settings as a constraint (Garcia et al., 2005; Moro et al., 2002). The biphasic current waveform (*P*_*BW*_) was generated as the following,

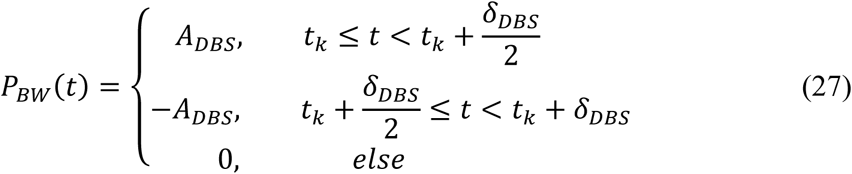

where, *t*_*k*_ are the onset times of the current pulses, *A*_*DBS*_ is the amplitude of the current pulse, *δ*_*DBS*_ is the current pulse width.

DBS stimulation with biphasic current waveform and four contact point configuration is given as,

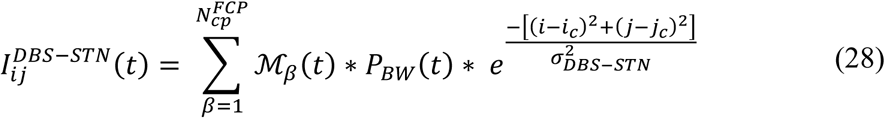

where, 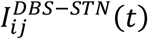 is the DBS current received by STN neuron at position (*i, j*) considering the lattice position (*i*_*c*_, *j*_*c*_) as the electrode contact point at time (*t*), ℳ_*β*_(*t*) is the indicator function which controls the activation of stimulation site *β*, 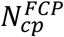 is the number of activated stimulation contact points 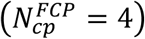, *P*_*BW*_(*t*) is the biphasic current waveform at time *t, σ*_*DBS*−*STN*_ is used to control the spread of stimulus current in STN network.

In the proposed excitotoxicity model, the DBS therapy was implemented by the following criterion,

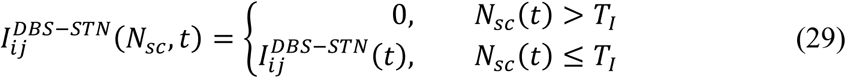

where, 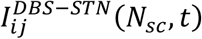 is the stimulation current to STN neuron at position (*i, j*) based on the number of SNc cells surviving at a particular time (*t*), *N*_*sc*_(*t*) is the number of SNc neurons surviving at a particular time (*t*), *T*_*l*_ represents the number of SNc cells surviving at which intervention of a particular treatment was employed.

DBS therapy in STN based on disruptive hypothesis proposed earlier (Muddapu et al., 2019; Rosenbaum et al., 2014) was implemented by the following criterion,

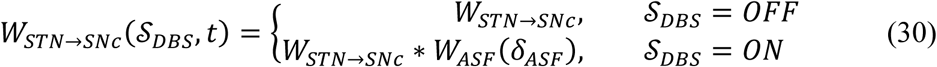

where, *W*_*STN*→*SNc*_(𝒮_*DBS*_, *t*) is the change in synaptic weight of STN to SNc based 𝒮_*DBS*_ = {*ON, OFF*} at a particular time (*t*), 𝒮_*DBS*_ is indicator function which modulate the activation and inactivation of DBS, *W*_*ASF*_ is the weight matrix based on the percentage of axonal and synaptic failures (*δ*_*ASF*_ = 5, 10, 20, 30, 40, 50, 60, 70, 80, 90, 100). The DBS parameters are given in Table-4.

**Table 4:** Parameter values of neuroprotective therapies.

| Parameter | Value | Parameter | Value |
| --- | --- | --- | --- |
| DBS frequency ($f_{DBS}$) in Hertz | 130 | Biphasic pulse width ($\delta_{DBS}$) in microseconds | 200 |
| Biphasic DBS amplitude ($A_{DBS}$) in picoampere | 1000 | Spread of the current ($\sigma_{DBS}$) | 5 |
| Population size of SNc ($P_{snc}$) | 64 | | |

#### Calcium Channel Blockers

It was reported that calcium channels in the SNc neuron contribute to neurodegeneration in PD (Benkert et al., 2019). In calcium channel blocker therapy (involving calcium blockers such as dihydropyridine, amlodipine), the excess calcium influx into SNc neurons is blocked by reducing the flux through calcium channels in the proposed model (Liss and Striessnig, 2019; Ritz et al., 2010). In the proposed excitotoxicity model, the calcium channel blocker therapy was implemented by the following criterion,

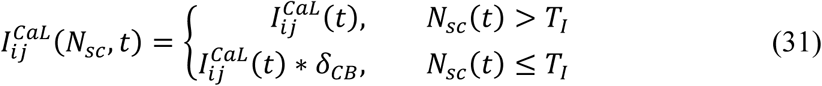

where, 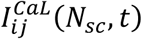 is the instantaneous calcium current of SNc neuron at position (*i, j*) based on the number of SNc cells surviving at a particular time (*t*), *N*_*sc*_(*t*) is the number of SNc neurons surviving at a particular time (*t*), *δ*_*CB*_ is the extent of calcium channel blockers, *T*_*l*_represents the number of SNc cells surviving at which intervention of a particular treatment was employed.

#### Enhancement of CBP Expression

It was reported that the expression of CBPs in general is very low in case of SNc neurons (Chung et al., 2005; Greene et al., 2005; Mendez et al., 2005) and reduces even further under PD (Fairless et al., 2019; Hurley et al., 2013; Yamada et al., 1990; Zaichick et al., 2017). Overexpression of CBPs was found to be neuroprotective in PD (Inoue et al., 2019; McLeary et al., 2019; Yuan et al., 2013). In the enhancement of CBPs expression, the calcium-binding proteins such as calbindin (Calb) and calmodulin (Cam) concentration was increased by the administration of external CBP (*δ*_*ECP*_). In the proposed excitotoxicity model, the enhancement of CBPs expression therapy was implemented by the following criterion,

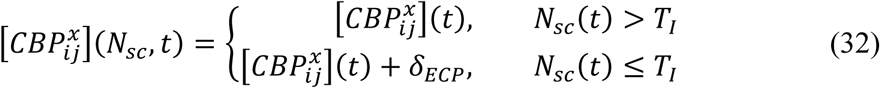

where, 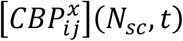 is the CBP concentration based on the number of SNc cells surviving at a particular time (*t*) (*N*_*sc*_(*t*)), 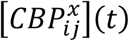 is the instantaneous CBP concentration where *x* = {*Calb, Cam*}, *δ*_*ECP*_ is the extent of CBP enhancement, *T*_*l*_ represents the number of SNc cells surviving at which intervention of a particular treatment was employed, *N*_*sc*_(*t*) is the number of SNc neurons surviving at a particular time (*t*).

#### Apoptotic Signal Blockers

It is known the neurons which are in stress (increase calcium levels) undergo neurodegeneration through apoptosis (Michel et al., 2016). In apoptotic signal blocker therapy (such as azilsartan, TCH346, and CEP-1347), the apoptotic signaling pathways are blocked by inhibiting the activation of caspase 9 and caspase 3 (Gao et al., 2017; Perier et al., 2012; Yacoubian and Standaert, 2009). In the proposed excitotoxicity model, the apoptotic signal blocker therapy was implemented by the following criterion,

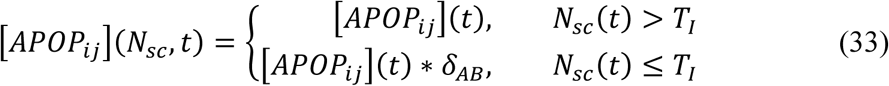

where, [*APOP*_*ij*_](*N*_*sc*_, *t*) is the apoptotic signal based on the number of SNc cells surviving at a particular time (*t*) (*N*_*sc*_(*t*)), [*APOP*_*ij*_](*t*) is the instantaneous apoptotic signal, *δ*_*AB*_ is the extent of apoptotic signal blockage, *T*_*l*_ represents the number of SNc cells surviving at which intervention of a particular treatment was employed, *N*_*sc*_(*t*) is the number of SNc neurons surviving at a particular time (*t*).

## RESULTS

We have investigated the neuronal model of SNc for their characteristic firing patterns (Figure 2) and studied the effect of energy deficiency on the model response (Figure 3). We then extensively studied the effect of energy deficiency and overstimulation by STN on the survivability of SNc neurons (Figure 4). Finally, we have explored various therapeutics such as glutamate inhibition, dopamine restoration, subthalamotomy, deep brain stimulation, calcium channel blockers, enhancement of CBPs, and apoptotic signal blockers (Figure 5,6) which slow down the SNc cell loss resulting in neuroprotective.

**Figure 2:**
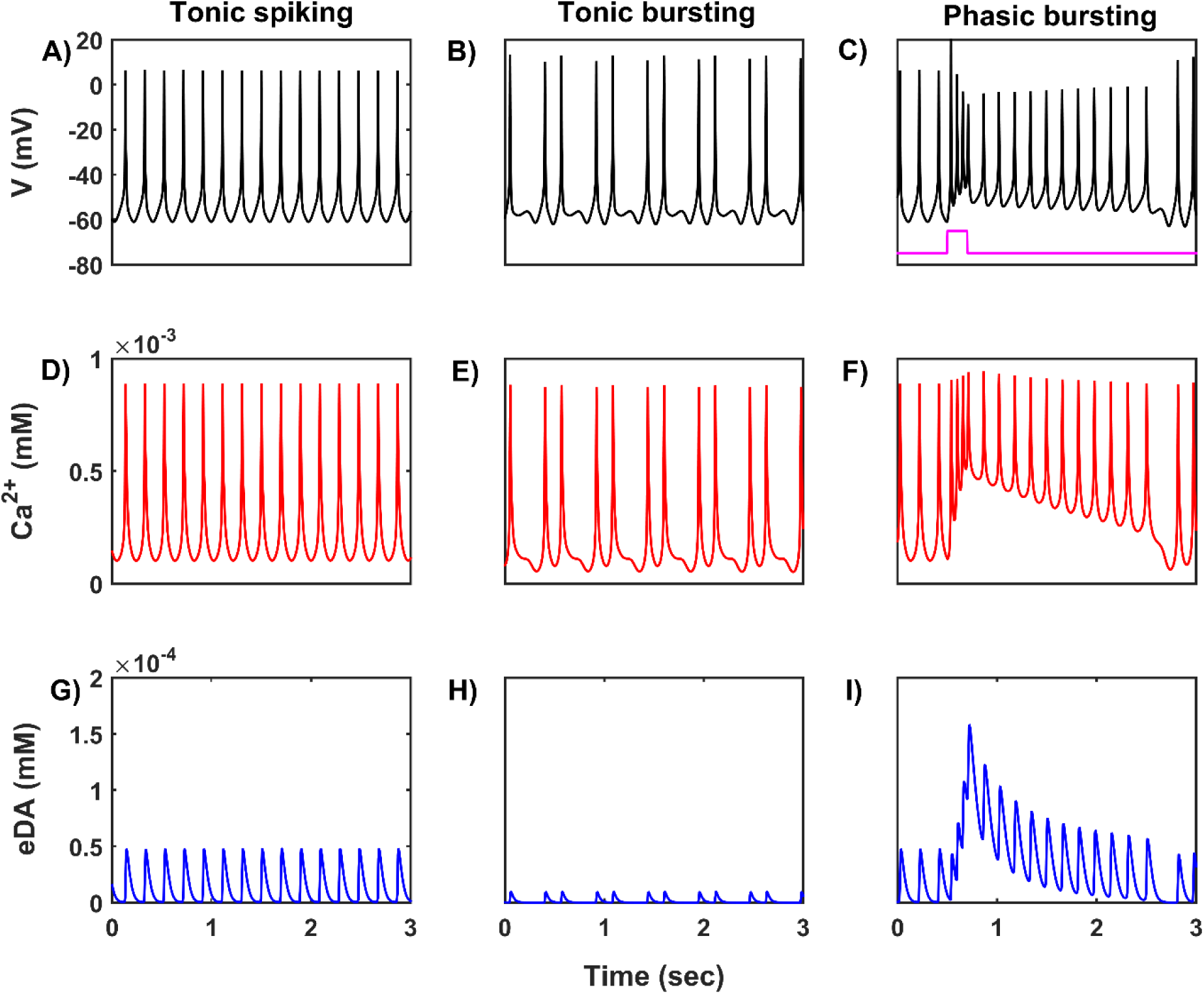
Characteristic firing patterns of the SNc neuronal model. Oscillations of membrane voltage potential **(A, B, C)**, intracellular calcium concentration **(D, E, F)**, and extracellular dopamine concentration **(G, H, I)** in tonic spiking, tonic bursting and phasic bursting. SNc, substantia nigra pars compacta; V, voltage; Ca^2+^, calcium; eDA, extracellular dopamine; mV, millivolt; mM, millimolar; sec, second.

**Figure 3:**
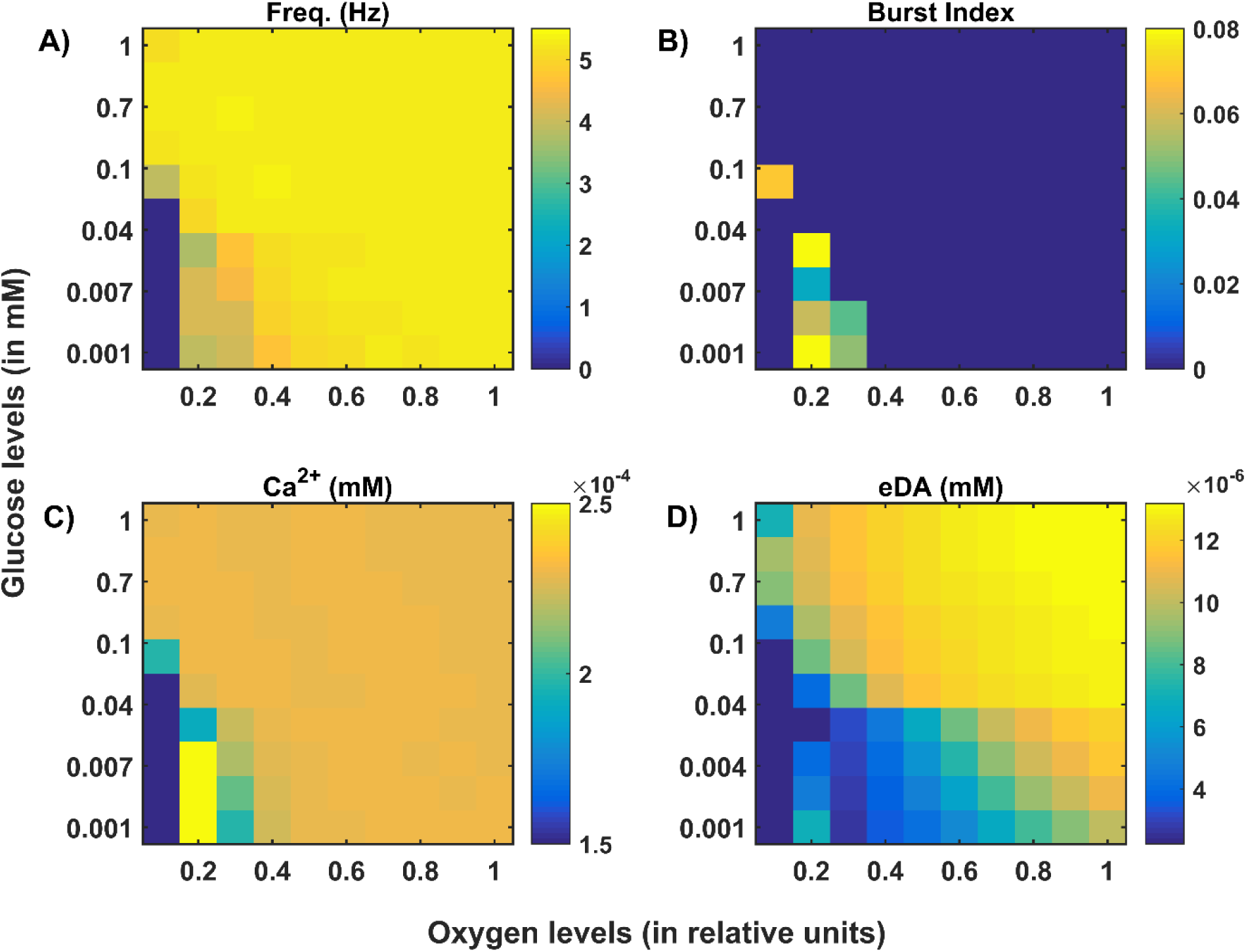
Model response of SNc neurons under hypoglycemia and hypoxia conditions. **(A)** Average frequency of firing. **(B)** Burst index. **(C)** Intracellular calcium concentration. **(D)** Extracellular dopamine concentration. SNc, substantia nigra pars compacta; Ca^2+^, calcium; eDA, extracellular dopamine; DA, dopamine; Hz, hertz; mM, millimolar.

**Figure 4:**
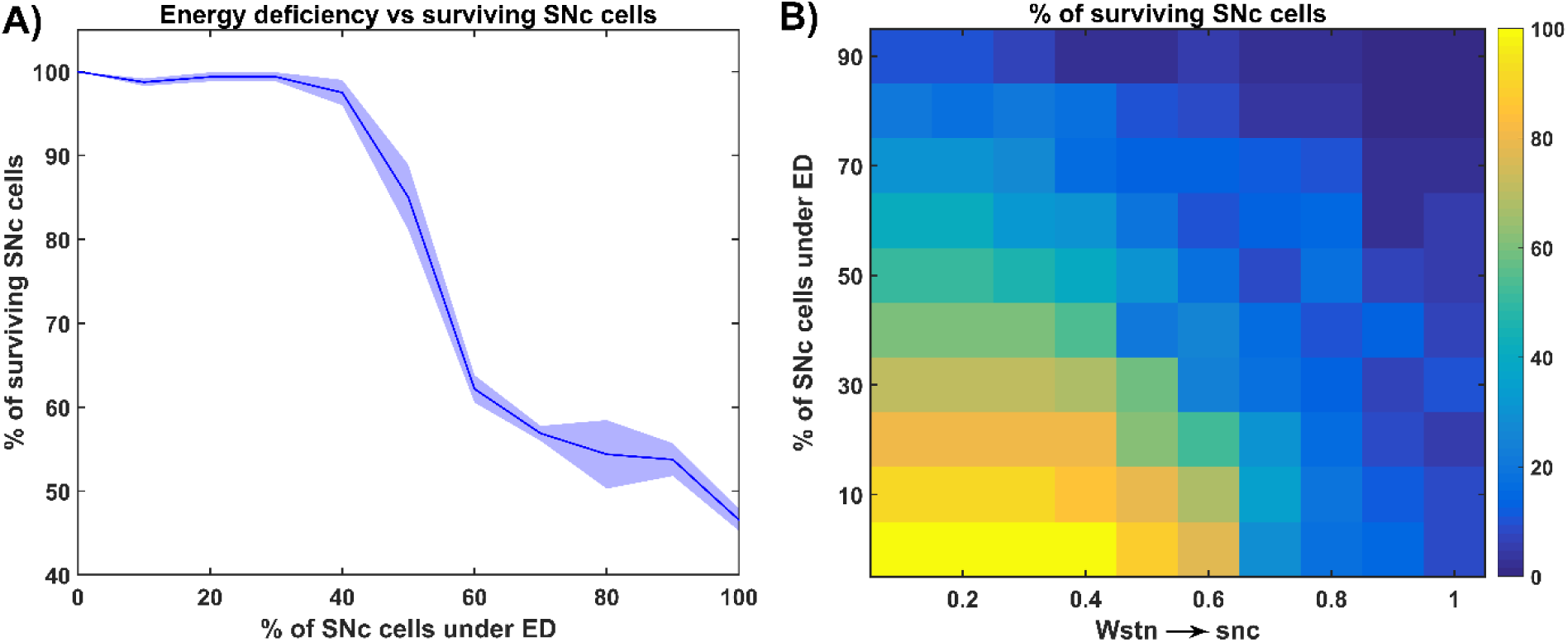
Response of the proposed hybrid model of excitotoxicity under energy deficiency. **(A)** Percentage of surviving SNc cells against the percentage of SNc cells under ED. **(B)** Percentage of surviving SNc cells for varying percentage of SNc cells under ED and synaptic weight from STN to SNc. STN, subthalamic nucleus; SNc, substantia nigra pars compacta; ED, energy deficiency.

**Figure 5:**
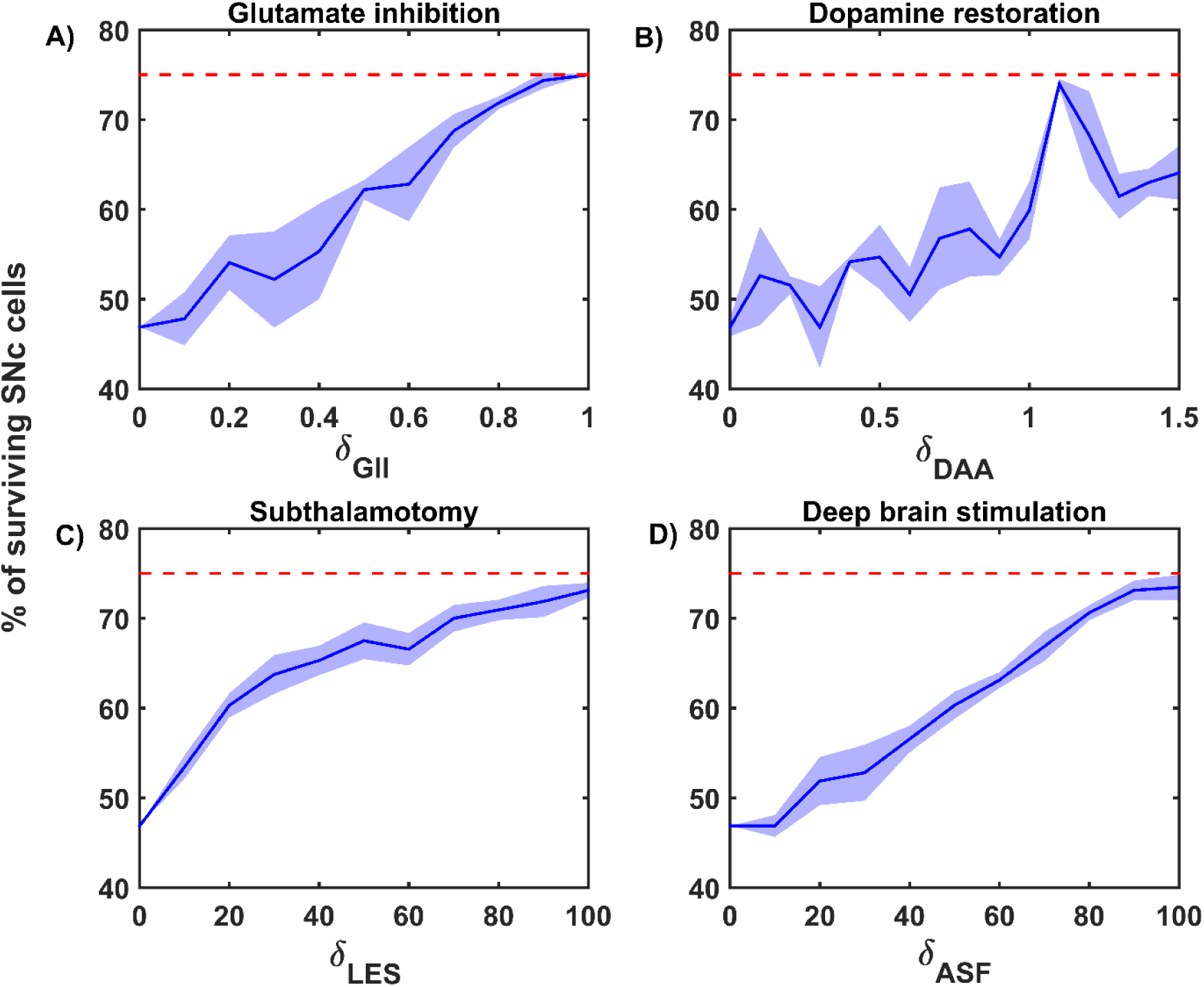
Proposed model response for different neuroprotective therapies. Percentage of surviving SNc cells in glutamate inhibition **(A)**, dopamine restoration **(B)**, subthalamotomy **(C)**, and deep brain stimulation **(D)** therapies. Red trace indicates the percentage loss of SNc cells at which intervention of particular therapy was initiated. SNc, substantia nigra pars compacta; GI, glutamate inhibition; DR, dopamine restoration; LES, STN lesioning; ASF, axonal and synaptic failures.

**Figure 6:**
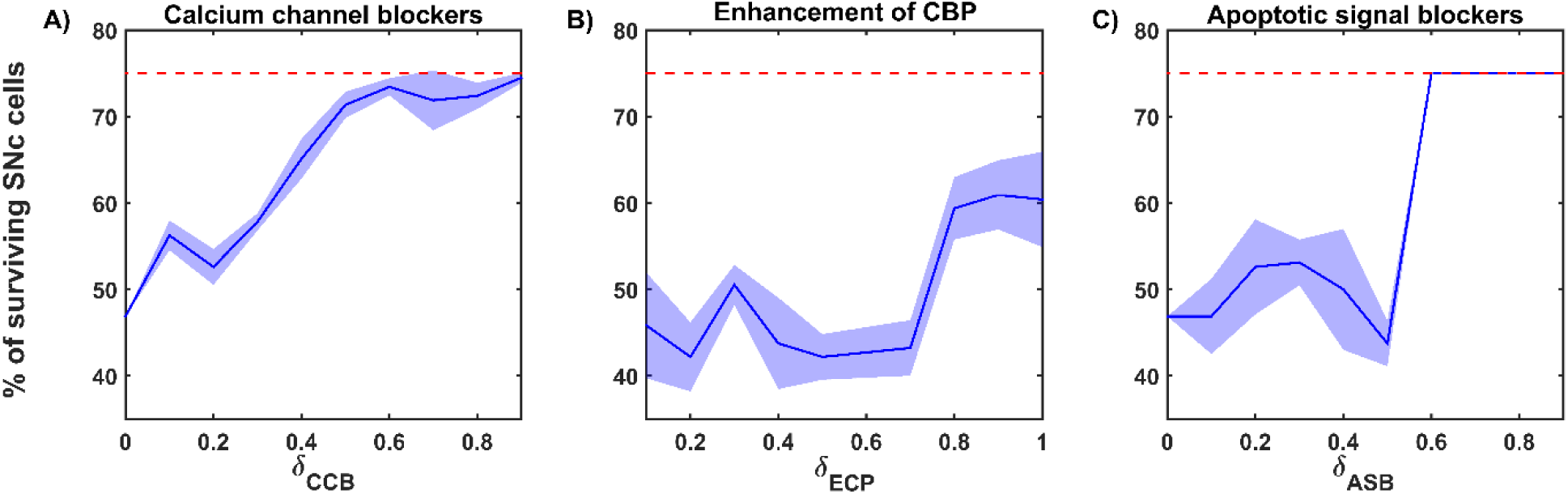
Proposed model response for different neuroprotective therapies. Percentage of surviving SNc cells in calcium channel blockers **(A)**, enhancement of calcium-binding proteins **(B)**, and apoptotic signal blockers **(C)** therapies. Red trace indicates the percentage loss of SNc cells at which intervention of particular therapy was initiated. SNc, substantia nigra pars compacta; CCB, calcium channel blockers; CBP, calcium-binding proteins; ECP, enhancement of calcium-binding proteins; ASB, apoptotic signal blockers.

### Characteristic Firing of SNc Neuron

SNc neurons exhibit two distinct firing patterns: low-frequency irregular tonic or background firing (3 − 8 *Hz*) and high-frequency regular phasic or bursting (∼ 20 *Hz*) (Grace and Bunney, 1984b, 1984a). In the proposed HEM model, the SNc neurons show spontaneous (tonic) spiking with a firing rate of ∼ 4 *Hz* (Figure 2A) (Grace and Bunney, 1984b). The intracellular calcium concentration of the SNc neuron during resting state was ∼ 1×10^−4^ *mM*, and it can rise to higher than 1×10^−3^ *mM* upon arrival of the action potential (Figure 2D) (Ben-Jonathan and Hnasko, 2001; Dedman and Kaetzel, 1997; Wojda et al., 2008). Dopamine released by SNc neuron during tonic spiking was peaked at ∼ 45 × 10^−6^ *mM*, which was in the range of (34 − 48) × 10^−6^ *mM* observed experimentally (Figure 2G) (Garris et al., 1997). Under energy deficiency condition (where intracellular ATP concentration was clamped at 0.411), SNc neurons exhibit tonic bursting with two spikes per burst in the absence of external current (Figure 2B) (Grace and Bunney, 1984a). The intracellular calcium concentration of the SNc neuron during resting state was below 1×10^−4^ *mM*, and rises up to 1×10^−3^ *mM* upon arrival of the action potential similar to tonic spiking (Figure 2E). Dopamine released by SNc neuron during tonic bursting, at its peak, was as low as ∼ 10 × 10^−6^ *mM* (Figure 2H), which was in the range of (7 − × 10^−6^ *mM* observed experimentally (Chen, 2005; Koshkina, 2006). When an external current is applied (continuous monophasic current with duration 200 *ms* and amplitude of 100 × 10^−6^ *pA*), SNc neuron exhibits phasic bursting with more than two spikes per burst (Figure 2C). The intracellular calcium concentration of the SNc neuron during resting state was ∼ 1×10^−4^ *mM*, and rises to a higher value than 1×10^−3^ *mM* upon stimulation (Figure 2F). Dopamine released by SNc neuron during phasic bursting peaks as high as ∼ 150 × 10^−6^ *mM* (Figure 2I) which is in the range of (150 − 400) × 10^−6^ *mM* observed experimentally (Schultz, 1998).

### The Effect of Energy Deficiency on SNc Neuron

Under energy deficient conditions, SNc neuron exhibits bursting with a decreased firing rate (Figure 3A, 3B). During bursting, the average intracellular calcium concentration of SNc neuron rise to a peak value of 2.5×10^−3^ *mM* (Figure 3C). Under energy deficient conditions, average extracellular dopamine concentration released by SNc neuron is very low (∼ 4×10^−6^ *mM*) compared to normal conditions (∼ 14×10^−6^ *mM*) (Figure 3D).

### The Effect of Energy Deficiency on Excitotoxicity in SNc

To understand the effect of energy deficiency on excitotoxicity in SNc, glucose, and oxygen levels are reduced to very low values. As the percentage of SNc cells in energy deficiency increases, the percentage of surviving SNc cells decreases in a sigmoidal manner (Figure 4A). When there are more than 40% of SNc cells under energy deficient conditions, there is a significant decrease in the percentage of surviving SNc cells.

### The Effect of Energy Deficit and STN on the SNc

In order to understand the influence of STN in SNc excitotoxicity under energy deficiency, glucose and oxygen levels were changed to very low values along with varying synaptic weight from STN to SNc (0 *to* 1 *with a step size of* 0.1). The percentage of surviving SNc cells, at low values of *W*_*stn*→*snc*_ (0.1 *and* 0.2), showed a linear decrease with an increase in the percentage of SNc cells under energy deficiency (Figure 4B). However, the pattern of the percentage of surviving SNc cells changes to sigmoidal at intermediate values of *W*_*stn*→*snc*_ (0.3 *to* 0.6) with an increase in the percentage of SNc cells under energy deficiency (Figure 4B). Beyond *W*_*stn*→*snc*_ values of 0.6, the percentage of surviving SNc cells was approaching zero, with an increase in the percentage of SNc cells under energy deficiency (Figure 4B).

### Neuroprotective Strategies

The proposed hybrid excitotoxicity model was extended to study the effect of different therapeutic interventions on the progression of SNc cell loss. The following therapeutic interventions were simulated: glutamate inhibition, dopamine restoration, subthalamotomy, deep brain stimulation, calcium channel blockers, enhancement of calcium-binding proteins, and apoptotic signal blockers. During the implementation of neuroprotective strategies, the percentage of SNc cells under energy deficiency was fixed at 80%, and synaptic weight from STN to SNc was fixed at 0.3.

#### Glutamate Inhibition Therapy

According to the protocol mentioned in the methods section, the neuroprotective effect of glutamate agonists and antagonists on the percentage loss of SNc cells was implemented. The glutamate inhibition therapy was initiated at 25% of SNc cell loss (indicated by red trace in Figure 5A) for varying extent of glutamate inhibition (*δ*_*GII*_). As the extent of glutamate inhibition increases, the percentage of surviving SNc cells increases in a linear fashion (Figure 5A). At higher values of *δ*_*GII*_, the percentage of surviving SNc cells was below 50%, and at lower values, it was around 75%, which was same as when the therapeutic intervention was initiated.

#### Dopamine Restoration Therapy

According to the protocol mentioned in the methods section, the neuroprotective effect of dopamine agonists on the percentage loss of SNc cells was implemented. The dopamine restoration therapy was initiated at 25% of SNc cell loss (indicated by red trace in Figure 5B) for varying extent of dopamine restoration (*δ*_*DAA*_). As the extent of dopamine restoration increases, the percentage of surviving SNc cells increases in a nonlinear fashion (Figure 5B). At lower values of *δ*_*DAA*_, the percentage of surviving SNc cells was below 50%. At higher values of *δ*_*DAA*_ (> 1.1), the percentage of surviving SNc cells was around 60%. However, at intermediate levels of *δ*_*DAA*_ (0.9 < *δ*_*DAA*_ ≤ 1.1), the percentage of surviving SNc cells peaks close to 75%.

#### Subthalamotomy

According to the protocol mentioned in the methods section, the neuroprotective effect of subthalamotomy on the percentage loss of SNc cells was implemented. The subthalamotomy therapy was initiated at 25% of SNc cell loss (indicated by red trace in Figure 5C) for varying extent of STN lesioning (*δ*_*LES*_). As the extent of subthalamotomy increases, the percentage of surviving SNc cells increases in a hyperbolic fashion (Figure 5C). At lower values of *δ*_*LES*_, the percentage of surviving SNc cells was below 50%. With just *δ*_*LES*_ value of 20%, the percentage of surviving SNc cells was increased to greater than 60%. For *δ*_*LES*_ values above 20%, the percentage of surviving SNc cells was steadily increased to a maximum value of 73% for *δ*_*LES*_ value of 100%.

#### Deep Brain Stimulation

According to the protocol mentioned in the methods section, the neuroprotective effect of DBS on the percentage loss of SNc cells was implemented. The DBS therapy was initiated at 25% of SNc cell loss (indicated by red trace in Figure 5D) for varying extent of axonal and synaptic failures (*δ*_*ASF*_). As the extent of axonal and synaptic failures increases, the percentage of surviving SNc cells increases in a linear fashion (Figure 5D). At lower values of *δ*_*ASF*_, the percentage of surviving SNc cells was below 50%, and at higher values, it peaked at 73% for *δ*_*ASF*_ value of 100%.

#### Calcium Channel Blockers

According to the protocol mentioned in the methods section, the neuroprotective effect of calcium channel blockers on the percentage loss of SNc cells was implemented. The calcium channel blocker therapy was initiated at 25% of SNc cell loss (indicated by red trace in Figure 6A) for varying extent of calcium channel blockers (*δ*_*CCB*_). As the extent of calcium channel blockers increases, the percentage of surviving SNc cells increases in a sigmoidal fashion (Figure 6A). At higher values of *δ*_*CCB*_, the percentage of surviving SNc cells was below 50%. At *δ*_*CCB*_ values of lesser than 0.5, the percentage of surviving SNc cells was increased to greater than 70%.

#### Enhancement of CBP

According to the protocol mentioned in the methods section, the neuroprotective effect of CBP enhancement on the percentage loss of SNc cells was implemented. The enhancing CBP therapy was initiated at 25% of SNc cell loss (indicated by red trace in Figure 6B) for varying extent of enhancement of CBP (*δ*_*ECP*_). As the extent of enhancement of CBP increases, the percentage of surviving SNc cells increases in a sigmoidal fashion (Figure 6B). At lower values of *δ*_*ECP*_, the percentage of surviving SNc cells was below 50%. At *δ*_*CCB*_ values of greater than 0.7, the percentage of surviving SNc cells suddenly increased to greater than 60%.

#### Apoptotic Signal Blockers

According to the protocol mentioned in the methods section, the neuroprotective effect of apoptotic signal blockers on the percentage loss of SNc cells was implemented. The apoptotic signal blockers therapy was initiated at 25% of SNc cell loss (indicated by red trace in Figure 6C) for varying extent of apoptotic signal blockers (*δ*_*ASB*_). As the extent of apoptotic signal blockers increases, the percentage of surviving SNc cells increases in a sigmoidal fashion (Figure 6C). At higher values of *δ*_*ASB*_, the percentage of surviving SNc cells was around 50%. At *δ*_*ASB*_ values of lower than 0.5, the percentage of surviving SNc cells seems to be same as when the therapeutic intervention was initiated. In order words, there is no SNc cell loss after therapeutic intervention was initiated.

## DISCUSSION

The goal of this work is to develop a hybrid model of excitotoxicity in SNc, which helps us in understanding the mechanism behind neurodegeneration due to excitotoxicity under energy deficiency conditions. The present model is the integration of the comprehensive SNc model, which we have developed earlier (Muddapu and Chakravarthy, 2020) and the previous model of excitotoxicity (Muddapu et al., 2019). From the simulation results, it suggests there exist compensatory mechanisms that try to maintain the calcium homeostasis even under energy deficiency conditions. We also suggest therapeutic intervention, which can be neuroprotective in nature. Moreover, the model also gives scope to explore more novel approaches to halt or slow down the SNc cell loss in PD.

From our previous computational studies (Chander and Chakravarthy, 2012; Chhabria and Chakravarthy, 2016), we were able to show that cortical neurons at lower ATP levels exhibit a bursting type of firing patterns. In the present model, we are able to show that SNc neurons exhibit a bursting type of firing pattern under low glucose and oxygen levels (Figure 3B). The firing rate of SNc neuron decreases under energy deficiency as a result of bursting, where bursting can be considered as a compensatory mechanism and also an indicator of energy imbalance (de Kloet et al., 2005; Sverrisdóttir et al., 1998; Sverrisdóttir et al., 2000). The intracellular calcium accumulation was a result of failed efflux of calcium by ATP-dependent calcium pumps due to energy deficiency. In the present model of SNc neuron, we have considered a calcium-induced dopamine release mechanism (Lee et al., 2009; Oheim et al., 2006). Even with higher levels of calcium, extracellular dopamine released by SNc neuron was low as a result of failed packing of dopamine into vesicles due to energy deficiency (Figure 3D) (Blakely and Edwards, 2012; Hnasko and Edwards, 2012).

The relationship between the percentage of SNc cells under energy deficiency and the percentage of surviving SNc cells is threshold-like, where there was a significant drop in the percentage of surviving SNc cells after more than 40% of SNc cells are under energy deficient conditions. It suggests that there might be mechanisms which compensate at low energy levels in order to enable normal functioning of SNc neuron (Navntoft and Dreyer, 2016).

Calcium plays an essential role in the normal functioning of a neuron; any imbalance in calcium homeostasis leads to pathogenesis (Orrenius et al., 2003). In the case of SNc neurons, these compensatory mechanisms may play an important role in maintaining calcium homeostasis. In the present SNc model, calcium fluctuations were monitored by three compensatory mechanisms: excess calcium binds to calcium-binding proteins (calbindin and calmodulin), excess calcium sequestered into ER by active pumps, and excess calcium is taken up into MT by mitochondrial calcium uniports. Under energy deficiency, calcium homeostasis is disrupted, resulting in calcium accumulation due to the inactivation of ATP-dependent calcium pumps.

As the first line of defense to maintain calcium homeostasis, excess calcium binds to locally available calcium-binding proteins in the cytoplasm. If calcium-binding proteins could not handle accumulated calcium, as the next line of defense, excess calcium is sequestered into ER by sarco/endoplasmic reticulum calcium-ATPase. If accumulated calcium exceeds the capacity of ER to sequester it, as a final line of defense, excess calcium is taken up by MT by mitochondrial calcium uniports. If accumulated calcium goes beyond the capacity of these three compensatory mechanisms, calcium builds up in both ER and MT, leading to neurodegeneration by ER-stress and mitochondrial-induced apoptotic mechanisms respectively (Figure 7) (Bano and Ankarcrona, 2018). Thus, the threshold-like relationship between the percentage of SNc cells under energy deficiency and the percentage of surviving SNc cells can be explained.

**Figure 7:**
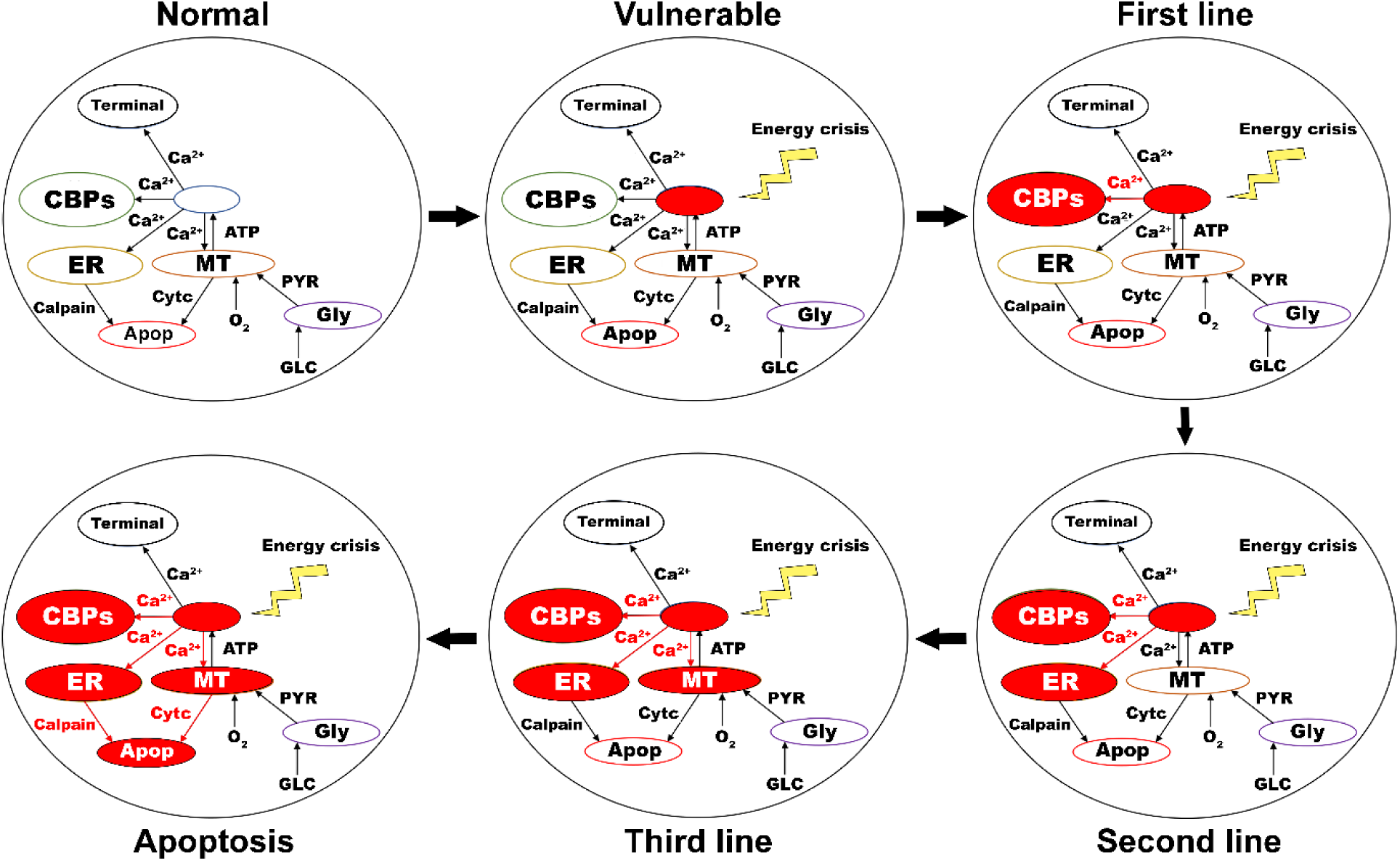
Proposed putative mechanism of excitotoxicity in SNc. Normal – healthy SNc neuron (normal calcium homeostasis); Vulnerable – vulnerable SNc neuron (compromise calcium homeostasis); First line – First line of defence to counter calcium imbalance by CBP (excess calcium binds to CBP); Second line – Second line of defence to counter calcium imbalance by ER (taken up into ER by SERCA); Third line – Third line of defence to counter calcium imbalance by MT (taken up into MT by MCUs); Apoptosis – ER-stress and mitochondrial-induced apoptotic mechanism activated (Calcium-induced neurodegeneration). SNc, substantia nigra pars compacta; ER, endoplasmic reticulum; MT, mitochondria; Ca^2+^, calcium; Cytc, cytochrome c; GLC, glucose; O_2_, oxygen; PYR, pyruvate; ATP, adenosine triphosphate; CBP, calcium-binding proteins; SERCA, sarco/endoplasmic reticulum Ca^2+^-ATPase; MCU, mitochondrial calcium uniports.

### Neuroprotective strategies

The reason behind the lower percentage of surviving SNc cells at higher values of *δ*_*GI*_ might be due to a mechanism known as ‘weak excitotoxicity’ (Albin and Greenamyre, 1992). Due to this mechanism, lower levels of available energy lead to calcium accumulation in SNc neurons, which undergo degeneration due to excitotoxicity. However, calcium accumulation in SNc neurons at lower values of *δ*_*GI*_ was low as a result of reduced excitation of STN on SNc even though they are exposed to energy deficiency. Thus, glutamate inhibition therapy proves to be neuroprotective to SNc neurons under energy deficiency conditions(Betts et al., 2012; Chan et al., 2010; Ferrigno et al., 2015; Masilamoni et al., 2011); moreover, it also seems to halt the SNc cell loss.

At intermediate levels of *δ*_*DAA*_ (0.9 < *δ*_*DAA*_ ≤ 1.1), the percentage of surviving SNc cells was maximum as a result of the total restoration of the dopaminergic drive to the STN neuronal population. However, beyond 1.1, *δ*_*DAA*_ results in lower percentage of surviving SNc cells when compared to intermediate levels. The reason behind the lower percentage of surviving SNc cells at higher values of *δ*_*DAA*_ might be due to the excessive activation of autoreceptors on the SNc neurons. There are four types of D2 autoreceptors on the SNc neurons, where they regulate neuronal activity and controls dopamine synthesis, release, and uptake (Ford, 2014). When these autoreceptors get activated, they result in reduced neuronal activity, dopamine synthesis, release, and uptake. In the present SNc model, we have considered all autoreceptors except one regulating dopamine uptake. At higher levels of dopamine, autoreceptors on the SNc neurons get overactivated, resulting in an overall decrease in dopamine release, which in turn reduced dopaminergic tone to STN neuronal population (Piccini and Pavese, 2006; Schapira and Olanow, 2003; Vaarmann et al., 2013). Decreased dopaminergic tone results in disinhibition of STN, which in turn leads to excitotoxic damage to SNc neurons (Rodriguez et al., 1998). Thus, dopamine restoration therapy proves to be neuroprotective to SNc neurons under energy deficiency. However, it can deteriorate the survivability of SNc neurons under high dosage (Lipski et al., 2011; Paoletti et al., 2019).

The relationship between the percentage of surviving SNc cells and *δ*_*LES*_ is threshold-like, where there was a significant rise in the percentage of surviving SNc cells with a *δ*_*LES*_ value of just 20%. The reason behind the lower percentage of surviving SNc cells at lower values of *δ*_*LES*_ might be due to a mechanism known as ‘strong excitotoxicity’ (Albin and Greenamyre, 1992). Due to this mechanism, overactivation of glutamatergic receptors leads to calcium accumulation in SNc neurons, which undergo degeneration due to excitotoxicity. It suggests that overexcitation from disinhibited STN is indeed a significant contributor to excitotoxic loss of SNc neurons (Figure 5C). However, *δ*_*LES*_value higher than 20%, the percentage of surviving SNc cells increases steadily with an increase in *δ*_*LES*_ values. At this point, excitotoxic loss of SNc neurons is due to ‘weak excitotoxicity,’ which is overcome by reducing excitatory drive from STN by subthalamotomy. Thus, subthalamotomy therapy proves to be neuroprotective to SNc neurons under energy deficiency, and in addition, it showed a substantial result with lower values of *δ*_*LES*_.

At lower values of *δ*_*ASF*_, the percentage of surviving SNc cells was below 50%. The reason behind the lower percentage of surviving SNc cells at lower values of *δ*_*ASF*_ might be due to ‘weak excitotoxicity’ (Albin and Greenamyre, 1992). As *δ*_*ASF*_ increases, the percentage of surviving SNc cells increases as a result of reduced excitation from disinhibited STN by DBS. We hypothesize that the neuroprotective effect of DBS therapy was as a result of STN pathological oscillations blockage from propagated to other nuclei; in order words increased axonal and synaptic failures of STN neurons due to DBS disrupts the information transfer through the stimulation site (Ledonne et al., 2012; Muddapu et al., 2019; Rosenbaum et al., 2014). Thus, DBS therapy appears to be neuroprotective to SNc neurons under energy deficiency, according to the proposed mechanism.

The relationship between the percentage of surviving SNc cells and *δ*_*CCB*_, *δ*_*ECP*_ and *δ*_*ASB*_ is threshold-like, where there was a sudden rise in the percentage of surviving SNc cells at higher values (Figure 6). The reason behind the lower percentage of surviving SNc cells at lower values of *δ*_*CCB*_, *δ*_*ECP*_ and *δ*_*ASB*_ might be due to ‘strong excitotoxicity’ (Albin and Greenamyre, 1992). It suggests that overexcitation from disinhibited STN is causing calcium accumulation in SNc neurons, which is overcome by reducing the influx of calcium into SNc neurons by blocking calcium channels at higher values of *δ*_*CCB*_ (Figure 6-A). Similarly, excess calcium accumulation in SNc neurons is overcome by enhancing CBP, which binds to free excess calcium and inactivates them at higher values of *δ*_*ECP*_ (Figure 6-B). It also suggests the neuroprotective effect of apoptotic signal blocker therapy at higher values of *δ*_*ASB*_ is a result of blockage of proapoptotic players (such as caspase 9, caspase 3) in SNc neuron which was activated due to excess calcium accumulation through ER-stress and mitochondrial-induced apoptotic mechanisms. Thus, calcium channel blocker, enhancement of CBP, and apoptotic signal blocker therapies prove to be neuroprotective to SNc neurons under energy deficiency; Moreover, apoptotic signal blocker therapy seems to halt the SNc cell loss.

In addition to the above mentioned therapeutic approaches, we have also simulated other therapeutic interventions such as mitochondrial calcium channel and permeability transition pore blockers, ATP supplements (phosphocreatine, creatine), and antioxidants which did not show neuroprotective effect in the present model. This can be due to the fact that direct ATP supplements and antioxidants might not be enough to show the neuroprotective effect in the present model.

### Limitations and Future Directions

Though the proposed model captures the exciting results of excitotoxicity, it is not without limitations. In the proposed model, the ischemic condition was implemented by modulating glucose and oxygen levels, which can be extended by adding a blood vessel module (Cloutier et al., 2009) and varying cerebral blood flow to simulate ischemia condition more realistically. In the proposed model, stress was monitored in SNc neurons alone, which can be extended to other neuronal types in the model by monitoring stress levels, where intracellular calcium accumulation can be a stress indicator (Bano and Ankarcrona, 2018). In order to do so, all neuronal types should be modeled as conductance-based models where calcium dynamics should be considered. The synaptic weights in the proposed model are not dynamic; we would like to incorporate learning principles such as spike time-dependent plasticity in the STN population, which can show the long-term effect of DBS treatment (Ebert et al., 2014; Iakymchuk et al., 2015). A more ambitious goal is to incorporate the present model into a large-scale model of basal ganglia (Muralidharan et al., 2018) in understanding the effect of therapeutics in terms of the behavioral response (Erdi et al., 2006). The future version of the model should be able to show the neuroprotective to other therapeutic interventions such as ATP supplements (phosphocreatine, creatine) and antioxidants.

### Conclusions

In conclusion, we believe that the proposed model provides significant insights in understanding the mechanism behind excitotoxicity in SNc neurons under energy deficiency conditions. From simulation results, it is evident that energy deficiencies occurring at cellular and network levels could precipitate the excitotoxic loss of SNc neurons in PD. From neuroprotective studies, it was clear that glutamate inhibition and apoptotic signal blocker therapies were able to halt the progression of SNc cell loss, a definitive improvement over other therapeutic interventions that only slow down the progression of SNc cell loss. Logically connecting with earlier our studies (Muddapu et al., 2019), we are presently reinforcing the idea of regarding PD as a metabolic disorder where metabolic deficiencies occurring at different levels in the neural hierarchy – subcellular, cellular, and network levels – precipitate the pathological changes of PD at multiple levels. Furthermore, using such models, we hope to be able to design and develop better therapeutics which target the root cause rather than the multitude of effects.

## CODE ACCESSIBILITY

The multi-scale excitotoxicity model code is available in ModelDB database (McDougal et al., 2017).

## AUTHOR CONTRIBUTIONS

VRM and VSC: conceived, developed the model, and prepared the manuscript.

